# Regression of juvenile tentacles is driven by loss of cell proliferation in *Haliclystus sanjuanensis*, a cnidarian with limited metamorphosis

**DOI:** 10.64898/2026.03.31.715438

**Authors:** Kennedy Bolstad, Leslie S. Babonis

## Abstract

Medusozoan cnidarians (*e.g.*, jellyfish) metamorphose from a benthic juvenile polyp into a pelagic adult medusa, providing a well-known example of a clade that uses tissue remodeling to create distinct juvenile and adult body plans. Staurozoans (*i.e.*, stalked jellyfish) are an atypical lineage of medusozoans that have lost their medusa stage; thus, their juvenile and adult body plans look remarkably alike. Their limited metamorphosis is characterized by the regression of primary (juvenile) tentacles and the development of secondary (adult) tentacles. In some staurozoan lineages, metamorphosis also involves development of novel adhesive structures (anchors), which are built on top of the regressing primary tentacles. Understanding how cells are partitioned from making juvenile tissues to making adult tissues is important for understanding how animals can make adult structures in the absence of complete metamorphosis. We compared the abundance and distribution of proliferative cells in tissues undergoing regression (primary tentacles) and development (secondary tentacles and anchors) during the juvenile to adult transition in the San Juan Island stalked jellyfish, *Haliclystus sanjuanensis*. We show that proliferative cells are lost in regressing primary tentacles but are gained in anchors, consistent with a shift in investment from juvenile to adult tissue. Prior to regression, primary and secondary tentacles show similar patterns in their proliferative cell distribution and in the identity of their cnidocytes (stinging cells), indicating that adult tentacles are made by re-deploying a juvenile tentacle program. Finally, we demonstrate that unlike secondary tentacles, primary tentacles cannot regenerate, illustrating that the temporary investment in this tissue is tied to their loss of proliferative cells. Thus, we propose that continued investment in a population of proliferating cells is an important mechanism for segregating temporary tissues (primary tentacles) from long-term tissues (secondary tentacles). These observations of cell dynamics in *H. sanjuanensis* suggest that temporary investment into juvenile structures may be used to pattern novel adult tissues, providing an important mechanism for diversifying adult body plans.

## Introduction

Examining the mechanisms by which diverse body plans arise is important for understanding the drivers of animal biodiversity. Tissue remodeling, or the reorganization of juvenile tissues into distinct adult structures, is a widespread mechanism across metazoans that facilitates the developmental segregation of juvenile and adult body plans during metamorphosis. These biphasic life cycles are thought to be advantageous for partitioning juvenile and adult phases into different niches, reducing competition for the same resources [1,2]. Notable examples include remodeling the larval esophagus during the dietary shift from planktivorous larvae to predatory adults in whelks [3] and remodeling the larval skeleton during the pelagic to benthic transition in sea urchins [4]. Outside of a few well-studied taxa [5,6], we still know very little about the dynamic cellular processes that drive the shift in investment from juvenile to adult body plans. Further, understanding whether juvenile tissues are transient, or whether they are important for building adult body plans is important for understanding the evolutionary mechanisms that drive metamorphosis.

Medusozoan cnidarians are a classic example of a clade with a biphasic life cycle segregated by tissue remodeling and the transition from benthic (polyp) to pelagic (medusa) niches. This clade is composed of hydrozoans (hydromedusae and hydroids), scyphozoans (true jellyfish), cubozoans (box jellyfish), and staurozoans (stalked jellyfish) (Fig 1A) [7]. Despite sharing the polyp-to-medusa transition, the mechanisms of tissue remodeling that drive this biphasic life cycle are distinct across medusozoans. Hydroids produce medusae by remodeling the polyp through lateral budding (Fig 1B) [8]. Scyphozoan medusae develop by remodeling the oral end of the polyp in a process called ‘strobilation’, where polyp tissues are segmented into one or more juvenile medusae (Fig 1C) [reviewed by 9]. Cubozoan medusae typically develop by remodeling the entire polyp (Fig 1D) [10,11], though some lineages only remodel their oral end like scyphozoans [12]. Staurozoans represent an unusual lineage of medusozoans that completely lack the pelagic medusa stage [13]. Considering the biphasic life cycle is thought to be ancestral to medusozoans [14], studying how the medusa stage was lost in staurozoans provides an interesting opportunity to examine how the loss of a life stage might lead to a new lineage of animals [15,16].

**Fig 1.**
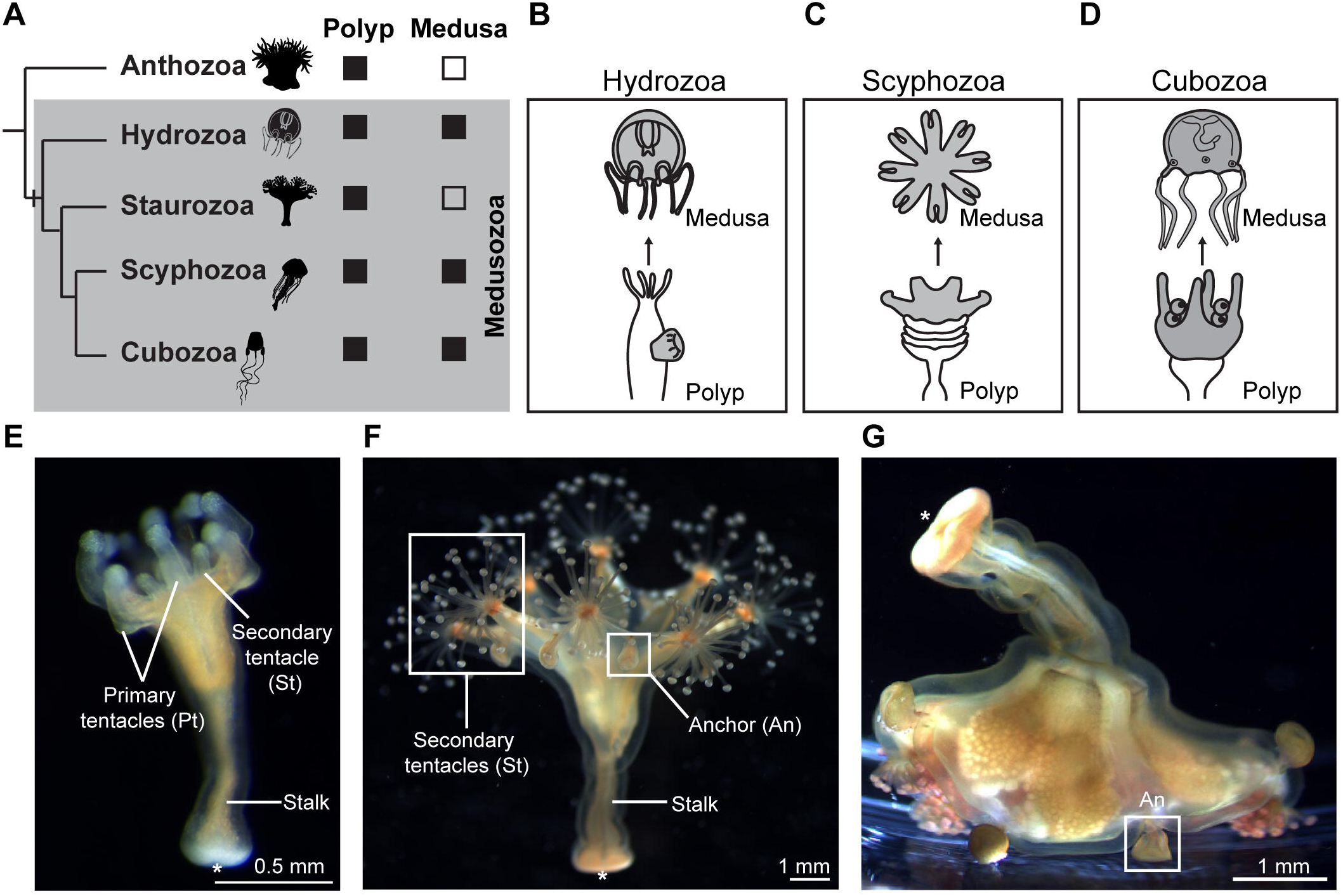
Staurozoans are medusozoans that lack a medusa stage. (A) Phylogeny of cnidarians showing which classes remodel juvenile polyps into adult medusa. The grey box highlights medusozoans and the hash mark indicates where metamorphosis from polyp to medusa was gained. Icons reproduced from PhyloPic (https://www.phylopic.org). (B-D) Polyp to medusa tissue remodeling across medusozoans. (B) Hydrozoan medusa budding off the body wall of a polyp. (C) Scyphozoan polyp undergoing strobilation to form juvenile medusae. (D) Cubozoan polyp transforming the oral end into a medusa. (E) Juvenile staurozoan with secondary tentacles (St) developing between primary tentacles (Pt). The base of the stalk is indicated with an (*). (F) Adult staurozoan with clusters of secondary tentacles found between anchors (An). (G) Adult staurozoan using anchors to adhere to a glass dish.

Because they lack a pelagic medusa stage, metamorphosis in staurozoans is highly reduced. Adult staurozoans develop medusa-like characteristics (e.g., gonads, gastric filaments, feeding tentacles) while retaining their polyp-like morphology and benthic lifestyle [reviewed by 17]. In certain staurozoan lineages, limited tissue remodeling starts in juvenile staurozoans with regression of their primary (juvenile) tentacles; these tentacles are either replaced by adhesive structures called anchors [17, 18] or are reabsorbed and not replaced [17, 19]. During this primary tentacle regression, clusters of secondary (adult) tentacles develop between the regressing primary tentacles [17 and references therein]. These secondary tentacles are the primary prey capture tissue found in adult staurozoans; after their initial development, new tentacles are added to the clusters, which are maintained throughout adulthood. Understanding how decreased investment in juvenile structures (*i.e.*, primary tentacles) is regulated is important for learning how animals decouple juvenile from adult traits and the mechanisms that may be selected upon to drive the evolution of novel body plans.

*Haliclystus sanjuanensis* (Staurozoa: Haliclystidae) is an important lineage in which to examine tissue remodeling, as anchors are unique to staurozoans in the family Haliclystidae [17, 20]. This staurozoan forms small populations in the low intertidal to shallow subtidal on the Pacific coast of North America from Alaska to California [21]. *H. sanjuanensis* has eight primary tentacles (Fig 1E) that are remodeled into anchors, and eight secondary tentacle clusters (Fig 1E,F). Anchors, which are populated by gland cells [20, 22–23], are thought to allow for greater adhesion to algae in the intertidal and subtidal environments (Fig 1F,G). Within Haliclystidae, primary and secondary tentacles are thought to have similar function (*i.e.*, feeding), morphology, longitudinal musculature, and innervation by FMRFamide neurons [23, 24]. Given the unique life history of staurozoans, we can ask the following questions: (1) If the medusa stage is lost, how are adult structures (like feeding tentacles) made? (2) If metamorphosis is reduced, why retain juvenile structures instead of developing directly? In this study, we compared the cell proliferation and regenerative capacity of primary and secondary tentacles in *H. sanjuanensis* and show that the loss of proliferative cells in primary tentacles indicates a loss of investment in this tissue. Further, we demonstrate that secondary tentacles are re-deployed primary tentacles with maintained regenerative capacity. We use these results to speculate about the role of juvenile tissues in scaffolding adult tissues.

## Methods

### Animal collection and care

*Haliclystus sanjuanensis* was collected at low tide from two locations on San Juan Island, Washington (USA): Cattle Point (48.452131° N, 122.961750° W) (Mills *et al*., 2023) and Grandma’s Cove (48.459125° N, 123.023947° W). Collections were conducted between May and July, from 2023-2025. Different developmental stages were opportunistically collected, but stage I animals tended to be found on branching algae (*e.g.*, *Odonthalia sp.*) while older stages (III and IV) were often found on *Ulva sp.* and *Mazzella sp.* Stage II animals were found on both *Odonthalia sp*. and *Ulva sp*. Given that *H. sanjuanensis* is found within small, concentrated populations, care was taken to not remove too many individuals in one season. Animals were kept in mesh enclosures (fish breeding enclosures, Anxingo, Amazon) in sea water tables at Friday Habor Laboratories (WA) with constant water flow and algae from their collection sites. Due to the constant water flow from the inlet, sea water temperature fluctuated between 8-11°C over the experimental period (May-July) [25]. Animals were fed every 2 days with newly hatched *Artemia salina* (Brine Shrimp Direct, USA). Live animals were imaged using an Infinity camera (v.3) mounted on a dissecting microscope. To capture staurozoan somersaulting behaviour, videos were taken using the Infinity Analyze software (v.7.1) and screenshots were taken via VLC Media Player (v.3.0.23).

### Tissue collection and EdU labelling

We used a Click-iT® EdU (5-ethynyl-2’-deoxyuridine) imaging kit (Invitrogen, C10338 and C10340) to label proliferating cells and their daughter cells in live tissues. Animals were immobilized for 30 minutes using menthol crystals added to 20 ml of seawater. Stock concentrations of 5-ethynyl-2’-deoxyuridine (EdU) and unlabelled thymidine (T1895, Sigma, USA) were initially dissolved in nuclease-free water before dilution to 100 µM in menthol containing seawater. After immobilization, animals were incubated in EdU for 30 minutes at 25°C. Following EdU incubation, animals were either washed three times in clean menthol seawater and fixed immediately or, washed three times in 100 µM unlabelled thymidine diluted in menthol seawater and three times in clean seawater before returning animals to their sea tables for pulse-chase experiments..

Tissues were fixed using 4% paraformaldehyde (PFA) with 0.2% glutaraldehyde for 1 minute at 25°C and 4% PFA without glutaraldehyde for 1 hour at 4°C. After fixation, tissues were washed three times for 15 minutes each in phosphate buffered saline with 0.1% Tween 20 (PTw) at 25°C. For combined immunohistochemistry and EdU labeling, animals were stored in fresh PTw at 4°C prior to analysis. For long term storage, EdU-labelled tissues were washed once with nuclease-free water and twice with 100% methanol before being stored in fresh 100% methanol at -20°C. Tissues stored in methanol were rehydrated with progressive washes into PTw. Tissues were permeabilized in phosphate buffered saline with 0.5% Triton X-100 (PTx) for 20 minutes before being placed into the EdU reaction cocktail for 30 minutes in the dark at 25°C following the manufacturer’s protocol. Tissues were then washed three times for 15 minutes each in PTw to remove excess reaction mix and nuclei were counterstained with 1.43 µM DAPI for 30 minutes at 25°C.

### Cnidocyte characterization

To characterize the cnidocyte types found in the tentacles and anchors of *H. sanjuanensis*, tissues were fixed as described above and stored in PTw at 4°C. Tissues were then mounted in a drop of 80% glyercol and squashed under a coverslip to dissociate the cells. Cnidocytes were characterized based on the morphology of their cnidocyst, which were imaged on a compound microscope (Nikon Eclipse E800).

### Immunohistochemistry

Following EdU development, tissues were washed in 0.2% PTx with 0.1% BSA (PBT) three times for 20 minutes each and blocked in 5% normal goat serum diluted in PBT for 1 hour at 25°C. Tissues were incubated in anti-γ-tubulin antibody (T5192, Sigma, USA) for 20 hours at 4°C before being washed five times for 10 minutes each in 0.2% PTx. Incubation in secondary antibody was also done for 20 hours at 4°C. Nuclei were counterstained with 1.43 µM DAPI for 30 minutes at 25°C.

### EdU pulse-chase experiments on isolated secondary tentacles

Animals with abundant secondary tentacles (stage III; Fig 2G,K) were used for this experiment, as they were the most abundant during collection. Animals were immobilized and approximately half of the tentacles from each tentacle cluster were cut using microdissection scissors (91500-09, Fine Science Tools, USA) (n=2 clusters per individual, totaling ∼25 tentacles per individual). Isolated tentacles were placed into EdU for 30 minutes and were either fixed immediately (n=5) or were washed three times for five minutes each in 100 µM unlabelled thymidine diluted in fresh sea water. Following EdU incubation and washing, isolated tentacles were placed into fresh seawater in a 6-well plate (29442-036, Corning) and maintained in the dark. To test the effects of cnidocyte firing on the number and distribution of proliferating cells, isolated tentacles were assigned to one of two groups (1) treatment with seawater only (control group, n=5 tentacles per individual), or (2) treatments with seawater containing live *Artemia* (to induce cnidocyte firing) (n=5 tentacles per individual). After the 30 minute treatment, *Artemia* were gently removed from the tentacles by gentle pipetting and tentacles were then placed into fresh seawater or seawater containing HU 24 well plate (29442-044, Corning), and maintained in the dark at 4°C to replicate cooler seawater conditions than room temperature. Seawater was replaced every 12 hours for the duration of the experiment. Isolated tentacles were collected at 48 hours post amputation (hpa) (n= 5 animals) or 96 hpa (n= 4 animals) and fixed for EdU labelling as described above.

**Fig 2.**
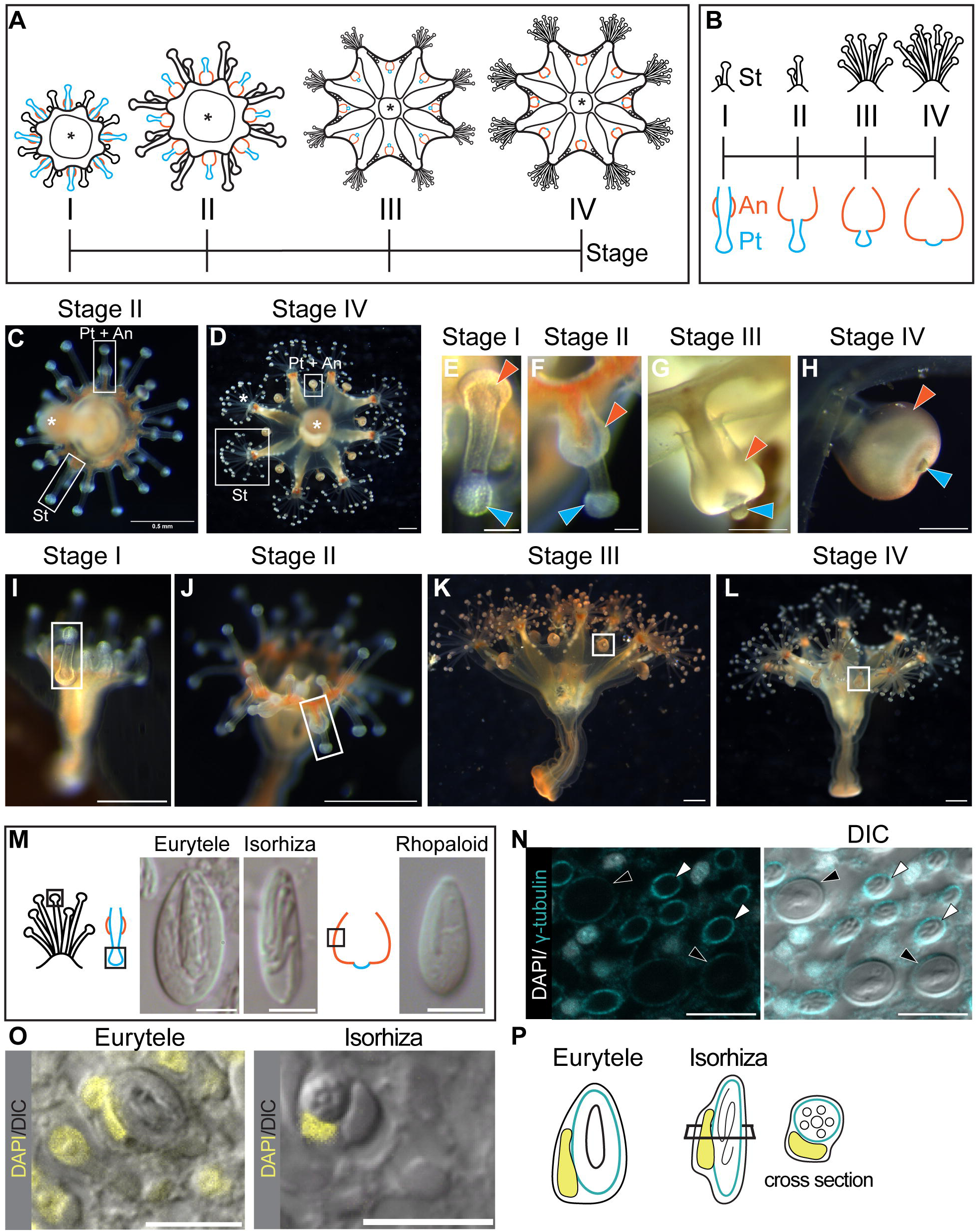
Staurozoans undergo limited metamorphosis and remodel primary tentacles into anchors. (A) Schematic staging guide for *H. sanjuanensis* representing an aboral view: eight secondary tentacle (St) clusters develop between eight regressing primary tentacles (Pt). Anchor (An) tissue develops on top of the regressing primary tentacles. The base of the stalk of this animal is indicated with an (*). (B) Schematic staging guide showing primary tentacle regression, anchor development, and secondary tentacle cluster growth in more detail. (C-D) Aboral view of live animals from stages II (C) and IV (D). (E-H) High magnification images showing different stages of the regressing primary tentacle (blue arrowhead) and developing anchor (orange arrowhead). (I-L) Whole animals, lateral view, during limited tissue remodeling. The regressing primary tentacles and developing anchors are indicated by white boxes. (M) Cnidocytes observed in the primary and secondary tentacles (microbasic euryteles and isorhizas) and anchor tissue (rhopaloids) are characterized by their cnidocyst pictured here with DIC microscopy. (N) Mature cnidocyte capsules labelled with anti-γ-tubulin antibody (cyan). Identity of the cnidocytes was verified using DIC (black arrowheads: euryteles, white arrowheads: isorhizas). (O) Morphology of the nuclei in mature cnidocytes (euryteles and isorhizas) labelled with DAPI. Mature cnidocytes are shown in DIC. (P) Cartoon showing mature cnidocyte nuclei morphology (yellow) and capsules (cyan). (N,O) Images are taken from a single slice of a confocal z-stack. Scale bars 0.1 mm (E, F), 0.5 mm (G-J), 1mm (K,L), 5 µm (M), 10 µm (N,O).

### Tentacle regeneration experiments

To explore the regenerative capacity of mature secondary tentacles, we followed the fate of tentacle clusters from which tentacles were removed for the experiments described above. Cut tentacle clusters (n= 2, per individual) and uncut tentacle clusters (n= 6, per individual) were imaged every 24 hours until 168 hpa when the animals were fixed. Tentacles were removed at 0hpa and then whole animals were treated with a 30 minute pulse of EdU at either 48 hpa (n= 5 animals) or 72 hpa (n= 4 animals). At these timepoints, one tentacle cluster per individual was removed and fixed, while the rest of the animal was washed with unlabelled thymidine to remove excess EdU and collected at 168hpa. Three additional animals (regressed primary tentacles, stage IV; Fig 2D,H,L) were opportunistically pulsed with EdU at 24hpa and collected immediately.

To compare the regenerative capacity of adult and juvenile tentacles, juveniles with prominent primary tentacles (stage II; Fig 2C,F,J) were used. Primary tentacles were cut partway down the tentacle stalk (n= 3 tentacles per individual, n= 3 animals total) and compared to unmanipulated primary tentacles (n=5 per individual) on the same individual. Animals were treated with a 30 minute pulse of EdU at 48hpa, washed with unlabelled thymidine to remove excess EdU, and collected at 168 hpa, as described above for the secondary tentacle regeneration experiments.

### Image collection, cell counting, and statistical analysis

Tissues were mounted in 80% glycerol and imaged by confocal microscopy on a Zeiss LSM 900 (Cornell University) or a Nikon C1Si (Friday Habor Lab, WA). Cell proliferation was quantified in 3D projected images produced from confocal z-stacks using Imaris software (Oxford Instruments) at the Cornell Institute of Biotechnology. The number of EdU-labelled cells within a tissue was counted and compared to the proportion of total nuclei. Statistical analyses were performed using R (v.4.5.1) and Rstudio (2025.05.1). Kruskal-Wallis tests were used to evaluate significant differences across groups and post hoc Dunn tests were used to evaluate which groups differed from each other.

## Results

To examine the cell dynamics of primary and secondary tentacles, we developed a staging guide for *H. sanjuanensis* (Fig 2A,B) and examined the cnidocyte (*i.e.*, stinging cell) composition of the two tentacle types and developing anchors; experiments throughout this paper will reference this guide. At the start of the transition from juvenile to adult, *H. sanjuanensis* has eight prominent primary tentacles and short secondary tentacles, with 1-3 secondary tentacles per cluster (Stage I; Fig 2E,I). Near the base of the primary tentacles are small bumps of tissue, marking the start of anchor development (Westlake and Page, 2017). At this stage, the primary and secondary tentacles are oriented in the same direction (orally) (Fig 2I). Stage II animals retain a long primary tentacle stalk with round anchor tissue extending towards the tentacle tip (Fig 2C,F,J). At this stage, primary tentacles begin to drop and orient opposite to the secondary tentacles, which are now longer than the primary tentacles. By stage III, animals have regressed their primary tentacles, so that only the rounded tentacle tip protrudes from the anchor tissue, which has become bulbous (Fig 2G,K). The tentacle tip is the last part of the primary tentacle to regress [21, 23]. From stage III onwards, secondary tentacles form dense clusters. By stage IV, only part of the primary tentacle tip is visible above the developed anchor (Fig 2D,H,L).

Cnidocytes are densely packed within primary and secondary tentacle tips and are occasionally found along the tentacle stalks [21]. Primary and secondary tentacles cannot be distinguished by their cnidocyte types, both have microbasic euryteles and isorhizas (Fig 2M). Cnidocytes within the anchor were sparse and identified as rhopaloids, similar to what are found within the body wall of *H. sanjuanensis* (Fig 2M) [21]. Mature cnidocyte capsules (euryteles and isorhizas) are labelled by anti-γ-tubulin antibodies (Fig 2N). Nuclei associated with mature cnidocytes were identified by their crescent shape (Fig 2O,P) [26].

### Cell proliferation decreases in primary tentacles during the tentacle- to- anchor transition

Prominent primary tentacles (Stage I) have a dense collar of proliferative cells around the base of the tentacle tip and a few scattered proliferative cells throughout the rest of the tentacle. Proliferative cells are abundant around the periphery of the developing anchor at this stage (Fig 3A-D). In stage II, proliferative cells become restricted to a tight collar around the base of the primary tentacle tip and to the proximal end of the growing anchor (Fig 3E-G). As the primary tentacle regresses further (Stages III and IV) the EdU collar in the primary tentacle is substantially reduced to only a few cells while the population of EdU- labelled cells in the proximal anchor tissue is maintained or expanded (Fig 3H-M). Quantification of the percentage of EdU-labelled cells found in the tentacle tip during this tissue transition shows a significant reduction of proliferative cells between stages I and IV (5.8 ± 3.4% to 0.3 ± 0.2%; *p-value* = 0.005) and stages II and IV (2.8 ± 1.9% to 0.3 ± 0.2%; *p-value* = 0.03) (Fig 3N). These results suggest there is a shift in investment from primary tentacle to anchor tissue (Fig 3O).

**Fig 3.**
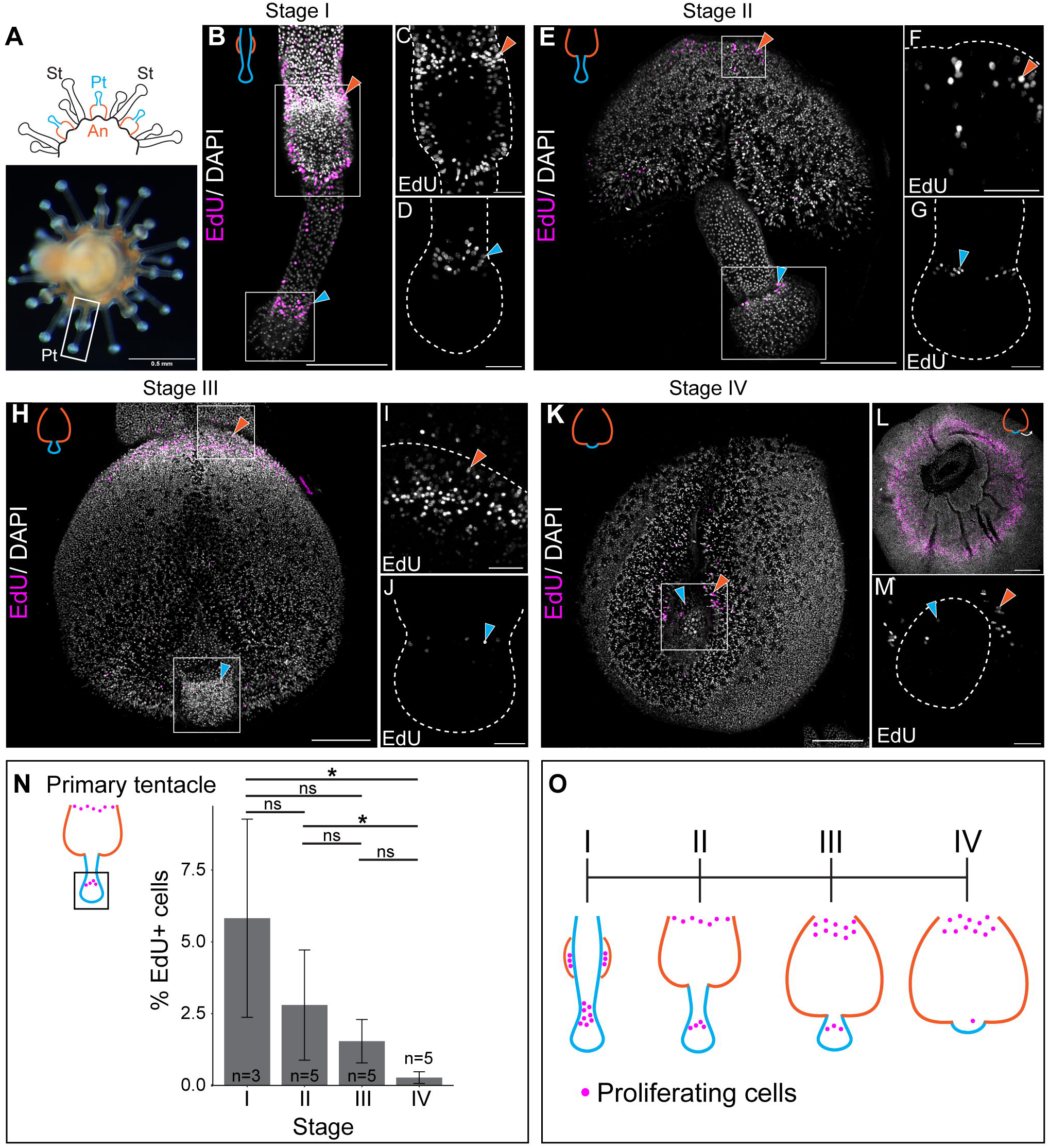
Cell proliferation during the transition from primary tentacle to anchor. (A) Primary tentacles (Pt) and growing anchors (An) sit between clusters of secondary tentacles (St). (B-M) The distribution of EdU- labelled nuclei (magenta) across regressing primary tentacle (D,G,J,M) and growing anchor tissue (C,F,I,) across stages I-IV. (L) EdU+ cells in an anchor, flipped over to show the portion of tissue that connects to the body wall. (N) Number of EdU- labelled nuclei in primary tentacle tips relative to total nuclei (DAPI). Significance tested with a Kruskal- Wallace test with post-hoc Dunn test, \**p* <0.05. N= 3 tentacles analyzed per individual of each stage, N= 3 individuals (Stage I) and N= 5 individuals (Stage II-IV). (O) Schematic summarizing the distribution of EdU- labelled cells in both tissues across stages. Scale bar 100 µm for low magnification images (B,E,H,K,L) and 25 µm for insets (C,D,F,G,I,J,M). Orange arrowheads point to proliferative cells in anchor tissue, blue arrowheads point to proliferative cells in primary tentacles. White dotted lines in the insets show the outline of the indicated tissue.

### Cell proliferation is maintained in secondary tentacles

Secondary tentacle clusters grow through continuous addition of more tentacles; thus, at any given life stage, these clusters will be populated by tentacles in many different stages of development (Fig 4A). At the onset of development, secondary tentacles first appear as a short stalk lacking the bulbous tentacle tip of established secondary tentacles; this bulbous tip is then developed prior to the elongation of the tentacle stalk (Fig 4A). These early tentacles are smaller than the existing primary tentacles and have proliferative cells throughout the stalk (Fig 4B,C,E,F), whereas established tentacles have a long stalk and proliferative cells restricted to a collar beneath the tentacle tip (Fig 4D,G,H). This pattern of cell proliferation in mature secondary tentacles is similar to the collar pattern found in Stage I primary tentacles (Fig 3B-D). To determine whether the reduced cell proliferation observed in regressing primary tentacles is also seen in mature secondary tentacles, we quantified EdU-labelled cells in established secondary tentacles in the same animals used to assay primary tentacle regression and found no significant difference in the abundance of proliferating cells in secondary tentacles across this time course (Fig 4I). Unlike what we have shown for primary tentacles, once proliferative cells become restricted to a collar beneath the tip of secondary tentacles, this proliferative cell population is maintained (Fig 4J). To identify the daughters of the EdU-labelled cells in the collar of secondary tentacles, we examined tentacles opportunistically labelled with EdU at the onset of development for a different pulse-chase experiment (Fig 4K). In early secondary tentacles with a bulbous tentacle tip (Fig 4C), both types of mature cnidocyte (*i.e.*, euryteles and isorhizas) were co-labelled with EdU and anti- γ-tubulin antibody (Fig 4L), confirming that at least some of the daughters of proliferative cells within the secondary tentacles are cnidocytes.

**Fig 4.**
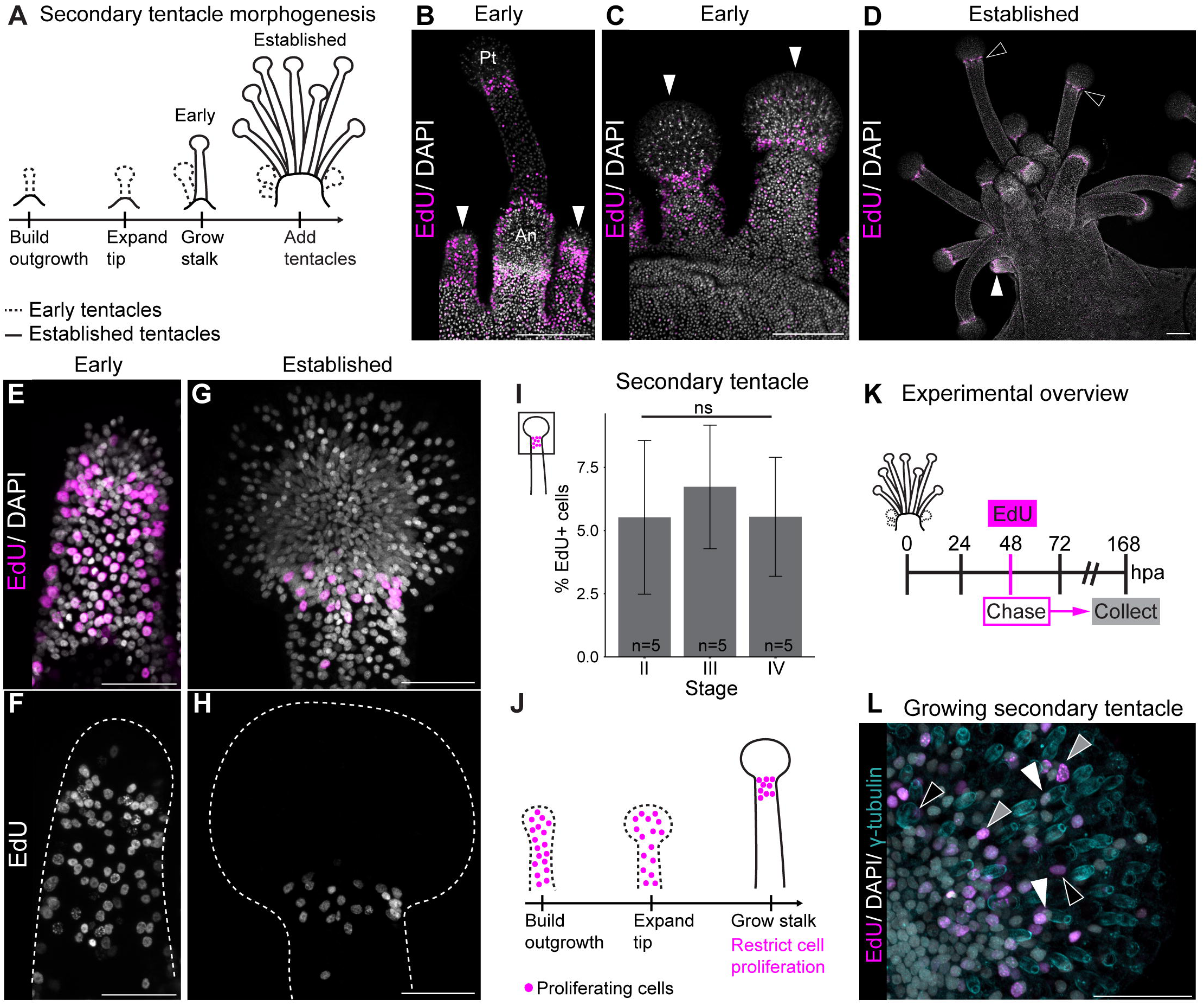
Cell proliferation during secondary tentacle growth and homeostasis. (A) Schematic showing growth and maintenance of secondary tentacles. (B, D) Early tentacles (white arrowheads) form as singlets in the space between primary tentacles (Pt) and developing anchors (An) and at the periphery of mature secondary tentacles (black arrowheads) in later stage animals. Growing secondary tentacles form a rounded tentacle tip prior to elongating the tentacle stalk. Distribution of EdU- labelled nuclei (magenta) across early (B,C,E,F) and established (D,G,H) secondary tentacle clusters. Images collected as follows: (B, E, F) Stage I, (G, H) Stage II, and (D) Stage III, correlating to stages shown in Fig 2. (I) Number of EdU- labelled nuclei relative to total nuclei in established secondary tentacle tips from animals in different stages. Kruskal-Wallace test, no significant differences observed. N= 3 tentacles analyzed per individual of each stage, N= 5 individuals per stage. (J) Schematic summarizing the distribution of EdU-labelled cells during tentacle morphogenesis. (K) Schematic outlining pulse chase experiments used for L. (L) Growing tentacle showing EdU- labelled euryteles (black arrowheads), isorhizas (white arrowheads) and non-cnidocyte cells (grey arrowheads). Scale bars 100 µm (B, C, D) and 25 µm (E-H, L).

### Proliferating cells move into the feeding tentacle tip over time but do not respond to cnidocyte discharge

Given the restricted distribution of proliferative cells within mature secondary tentacles, we hypothesize that cells within the collar are used to replace cells (*e.g.,* cnidocytes) within the tentacle tip. To directly test whether proliferative cells in the collar of secondary tentacles move into the tentacle tip in response to cnidocyte firing, or whether proliferative cells move into the tentacle tip at a constant rate, we conducted a pulse-chase experiment using EdU labeling in isolated, established secondary tentacles (Fig 5A). Cut tentacles were incubated in EdU and were either fixed immediately (0 hours post amputation (hpa)) or were chased in seawater with or without *Artemia* (Fig 5A-C). Since the rate of cell differentiation in staurozoans is currently unknown, we used to two chase periods: 48hpa and 96 hpa. We expected to see a difference in the abundance of proliferative cells between tentacles exposed to *Artemia* and those only treated with seawater, if cell proliferation is upregulated in response to cnidocyte firing. If cells are replaced at a constant rate within the tentacle, we expected that no difference would be observed (Fig 5D). There was no significant difference in the abundance of EdU-labelled cells, relative to total nuclei, in unfired (10.5 ± 9.2% 48 hpa; 6.4 ± 1.8% 96 hpa) and fired (10.6 ± 3.7% 48 hpa; 8.1 ± 2.9% 96 hpa) tentacles (Fig 5E,F). Proliferative cells from the collar moved into the tentacle tips of unfired controls (n= 3/5 animals chased until 48 hpa; n= 4/4 animals chased until 96 hpa) at the same rate as tentacles induced to fire with *Artemia* (n= 5/5 animals chased until 48 hpa; n= 4/4 animals chased until 96 hpa) (Fig 5G). Together, these results suggest that that cell proliferation is continuous in secondary tentacles and independent of cnidocyte firing.

**Fig 5.**
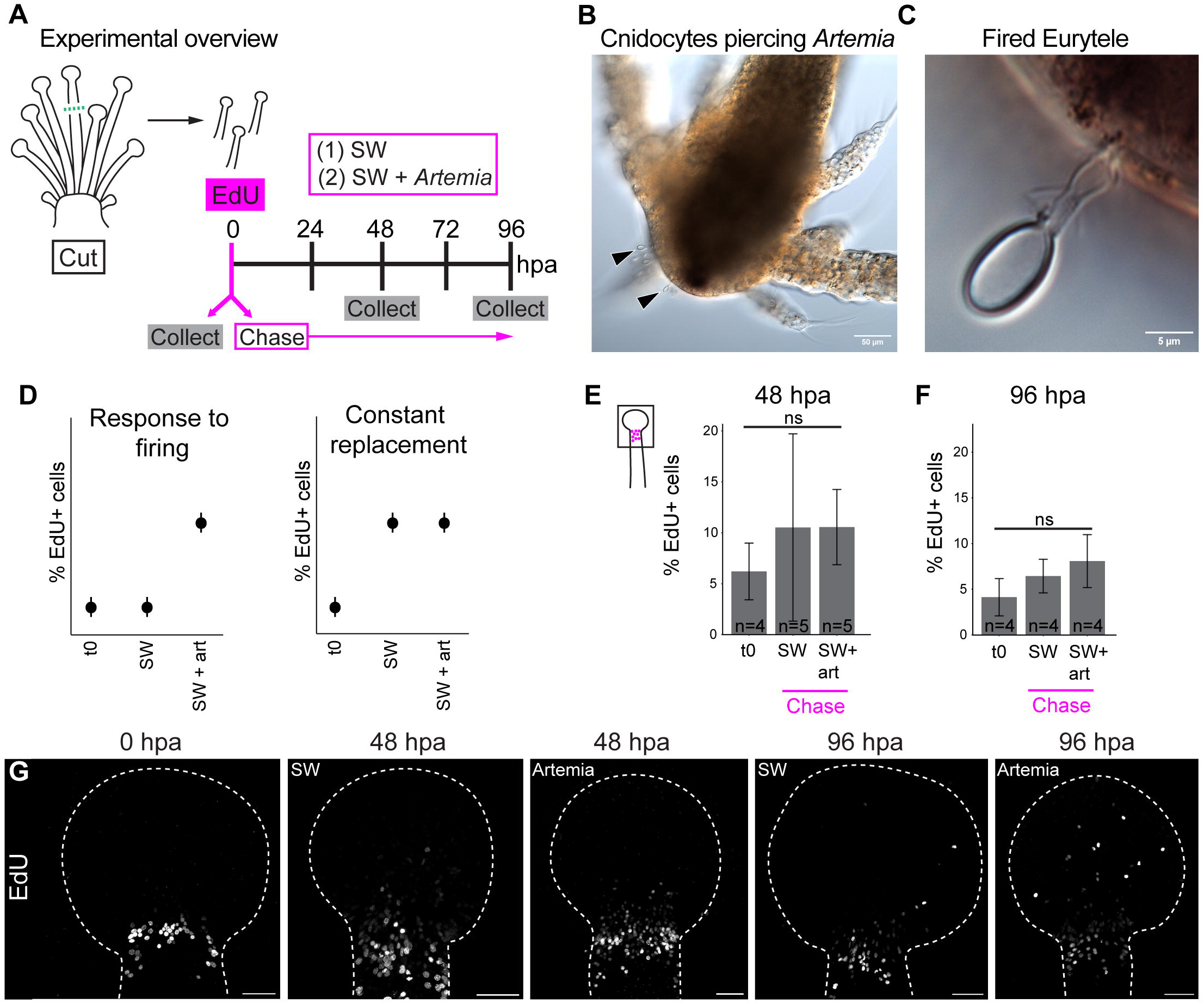
Proliferative cells move into the tip of secondary tentacles. (A) Schematic outlining the EdU pulse-chase experiments on cut secondary tentacles placed into seawater alone (SW) or seawater without *Artemia* (Art). (B) *Artemia* immobilized by fired cnidocytes (black arrowheads), (C) high magnification of a fired euryteles (cnidocyte) piercing the exoskeleton of *Artemia*. (D) Hypothesized outcomes for the experiments in (A): either EdU- labelled cells respond to firing (left), or cells replace cnidocytes at a constant rate (right), without a cue from fired cnidocytes. (E,F) Quantification of EdU- labelled cells in the cut tentacles. Kruskal-Wallace test with post-hoc Dunn test, no significant differences observed. N= 3 tentacles analyzed per individual, N= 5 individuals per treatment (48 hpa SW, 48 hpa SW+art), N= 4 individuals per treatment (48 hpa t0, all treatments 96 hpa). (G) Distribution of EdU- labelled cells in cut secondary tentacles across treatments. Scale bar 25µm. Hours post amputation (hpa). White dotted line shows outline of the tentacle.

### Secondary tentacles can regenerate but primary tentacles cannot

Given that staurozoans have proliferative cells restricted to a collar beneath their tentacle tips, we tested whether mature secondary tentacles cut mid-way down the tentacle stalk (below the collar of proliferative cells), could regenerate the rest of the tentacle (Fig 6). We compared regenerating tentacles to unmanipulated tentacles by cutting only a subset of mature tentacles from each cluster (Fig 6A-B). By 48 hpa, the cut tentacle stalks had healed, leading to a rounded tentacle stalk with some mature cnidocytes present (Fig 6C-E). By 72 hpa, the tentacle tip had thickened and started to become more pronounced than the tentacle stalk, with more mature cnidocytes present (Fig 6C-E). Secondary tentacles had fully replaced their tentacle tips by 168 hpa, however, they remained shorter than uncut tentacles (Fig 6C). This suggests that animals first invest in repopulating the mature cell types at the tip of the tentacle before investing in tentacle outgrowth during regeneration (Fig 6D). To examine the location of proliferative cells during regeneration, we incubated animals in EdU either at 48 hpa or 72 hpa and chased these until 168 hpa (Fig 6A). Uncut tentacles showed cell proliferation restricted to the base below the tentacle tip; at the same height as the cut site, uncut tentacles had only scattered proliferative cells (Fig 6F). Cut tentacles at 48 hpa had dense clusters of proliferative cells spanning from the tip of the healed tissue to partway down the tentacle stalk; at 72hpa dense clusters of proliferative cells were more restricted to the rounded tentacle tip (Fig 6F). At 48 hpa and 72 hpa proliferative cells were observed in both the ectoderm and the endoderm of cut tentacles, unlike the uncut tentacles (Fig 6E). At 168 hpa, the regenerated tentacle tip is filled with the daughters of proliferative cells and similarly to growing tentacles, EdU-labelled nuclei are associated both with mature cnidocytes and with non-cnidocyte cells (Fig 6F-G). Together, these results suggest that tentacle regeneration acts similarly to tentacle morphogenesis: once an initial stalk is in place, the tentacle tip is grown and mature cnidocytes are established prior to stalk elongation. Since there are few proliferative cells in the stalks of these tentacles, we suggest that one of three cell populations are contributing to this regeneration: (1) proliferative cells that travel from the elsewhere in the body, similarly to what is seen in hydrozoans, *C. hemisphaerica* and *Hydractinia echinata* (Hydrozoa) [27, 28], (2) quiescent cells in the stalk that are activated when the tissue is damaged, as seen in the medusa of *C. pacificum* (Hydrozoa) [26], or (3) de-differentiated epithelial cells used to repair damaged tissue, as hypothesized for non-hydrozoan cnidarians [29].

**Fig 6.**
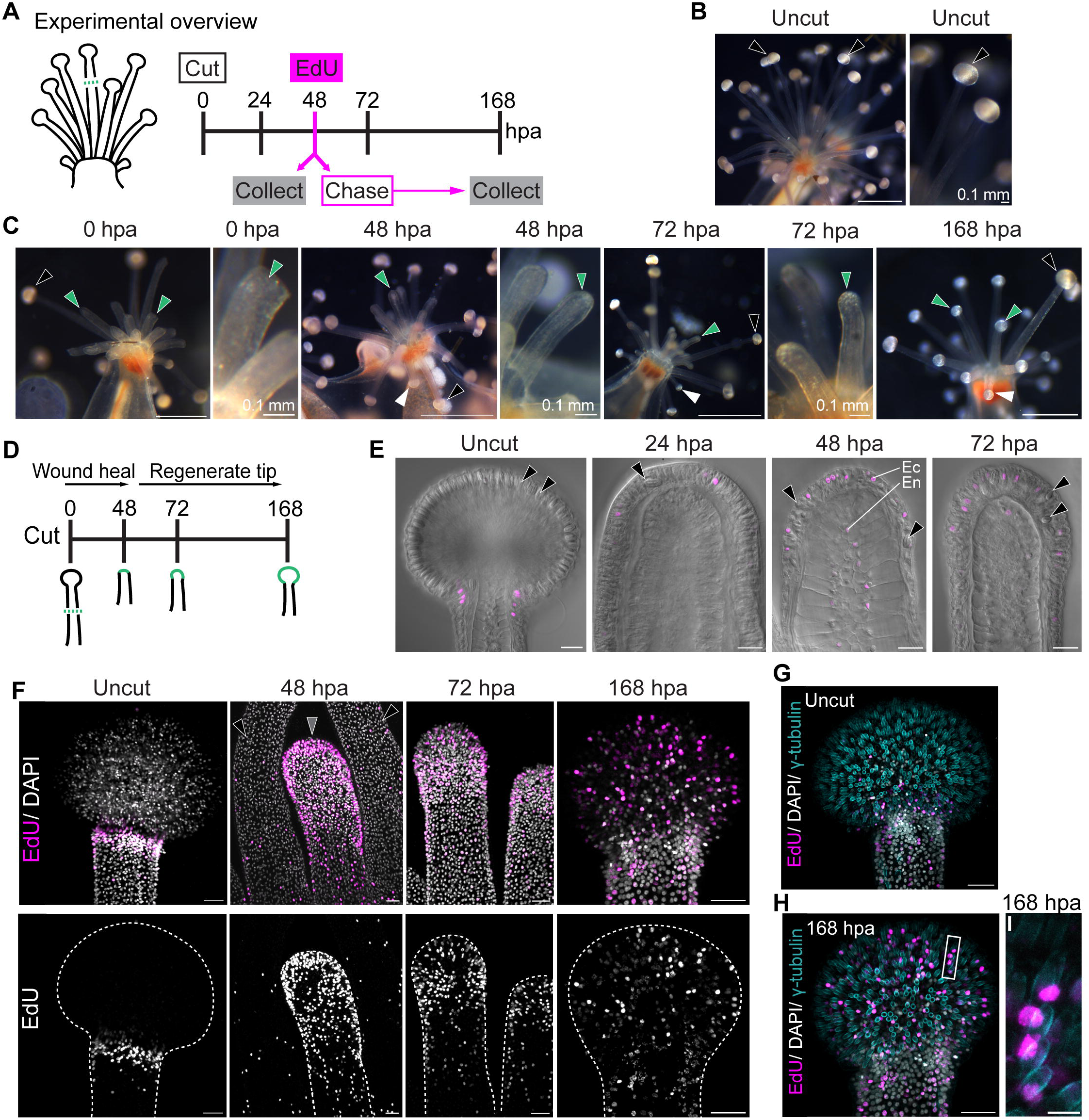
Secondary tentacles regenerate. (A) Schematic representation of the regeneration experiment. Tentacles were labelled with EdU either at 48 hours post amputation (hpa) or 72 hpa; only the 48 hpa pulse is shown. (B) Secondary tentacle clusters and a high magnification secondary tentacle in a live animal, prior to being cut. (C) Live images of representative secondary tentacle clusters during regeneration showing uncut tentacles (black arrowheads), regenerating tentacles (green arrowheads), and growing tentacles (white arrowheads). (D) Schematic summarizing secondary tentacle regeneration. (E) EdU- labelled nuclei (magenta) in uncut and regenerating tentacles; EdU- labelled nuclei are found in the ectoderm (Ec) and endoderm (En) at the tip of regenerating tentacles. Black arrowheads indicate mature cnidocytes (F) Top: EdU- labelled nuclei (magenta) in uncut and regenerating secondary tentacles. Nuclei counterstained with DAPI (white). Bottom: shows EdU-labelled nuclei only (white). (G-H) EdU-labelled cells move to the tentacle tips in regenerating secondary tentacles at 168 hpa (H), but largely remain in the collar in uncut tentacles (G). (I) Higher magnification inset of the tentacle shown in (H) showing EdU- labelled isorhizas. Scale bars 1mm (B, C), 20 µm (E), 25 µm (F-H), 5µm (I). (F-H) Are max projections from confocal z-stacks; (E) are from single optical sections.

Given the regenerative capacity of the secondary tentacles (Fig 6) and the decreased abundance of EdU-labelled in primary tentacles over time (Fig 3), we tested whether primary tentacles cut below the collar of proliferating cells could regenerate (Fig 7). These experiments were conducted in Stage II individuals (Fig 7A,B), where the collar of proliferative cells is obvious and anchor tissue has started to develop. Similar to regenerating secondary tentacles, the primary tentacles had healed the cut tissue by 48 hpa; however, the primary tentacle tips did not regenerate in the same time frame that secondary tentacle tips regenerated (168 hpa) (Fig 7C). Instead, the primary tentacle tissue was morphologically similar to the healed tissue at 48 hpa with very few proliferative cells at 168 hpa (Fig 7C-G). The abundant proliferative cells present in the anchors (Fig 7F-G) mirrored the proliferative cell abundance during the development of anchors from uncut primary tentacles (Fig 7D-E), suggesting that cell proliferation in the anchors was still behaving as expected. Together, these observations demonstrate differential regulation of regeneration in primary and secondary tentacles.

**Fig 7.**
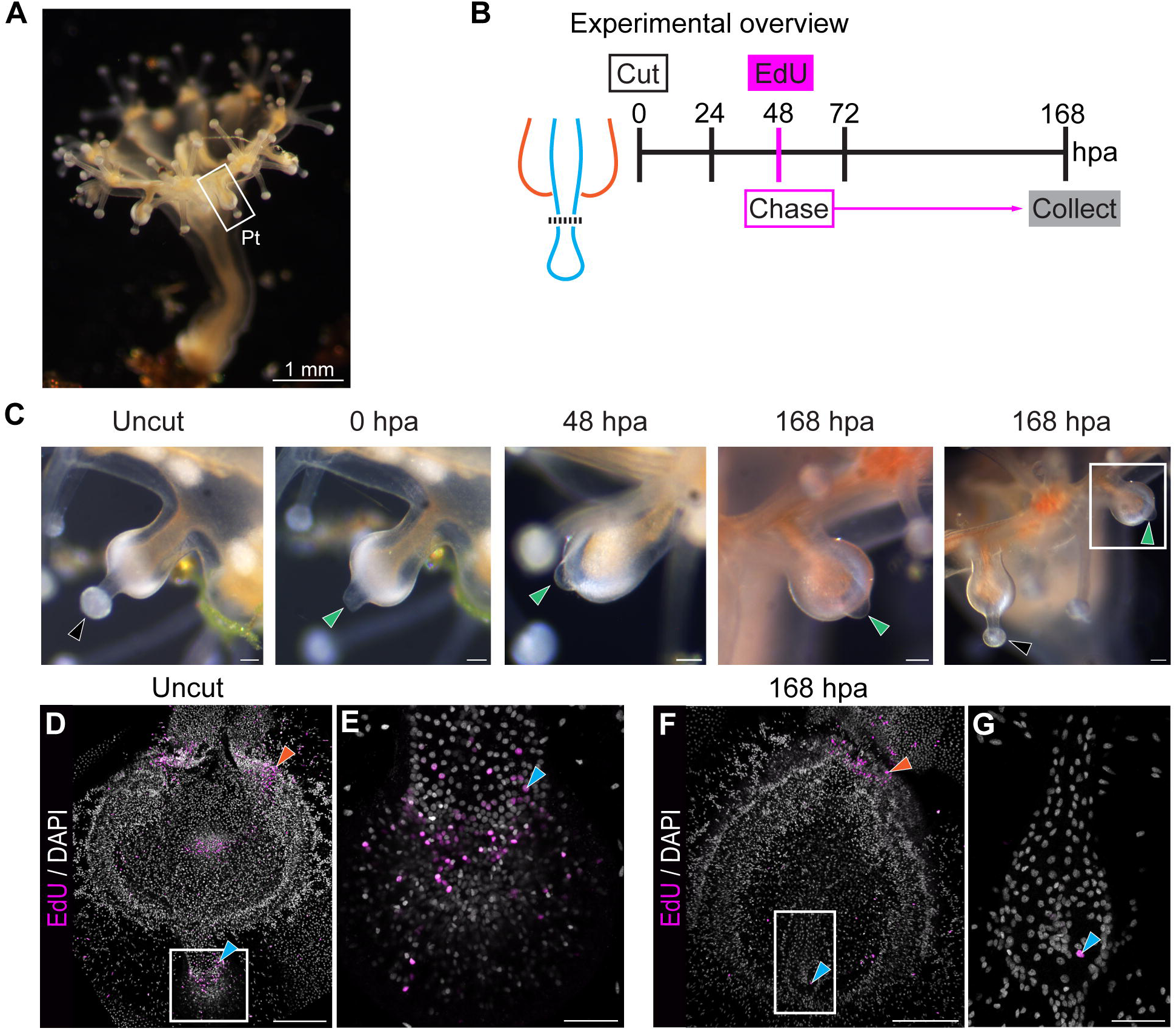
Primary tentacles do not regenerate. (A) Stage II animals were used for primary tentacle regeneration experiments. (B) Schematic of experiment. (C) Live images showing tissue morphology at the indicated times. Black arrowheads indicate uncut tissue and green arrowheads indicate cut primary tentacles. (D-G) Distribution of EdU- labelled nuclei (magenta) in uncut and cut tentacles. Nuclei were counterstained with DAPI (white). Boxes highlight regions highlighted in insets. Blue arrowheads indicate proliferative cells in primary tentacles, orange arrowheads point to proliferative cells in anchors. Scale bars 100 µm (C, D, F), 25 µm for insets (E, G).

## Discussion

Staurozoans are an understudied group of cnidarians; their highly reduced biphasic lifestyle makes them an important lineage for understanding the loss of metamorphosis. Using *H. sanjuanensis*, we have demonstrated different investment into primary and secondary tentacles: the degenerating primary tentacles reduce the number of proliferating cells and cannot replace amputated tissue (Figs 3,7), whereas secondary tentacles maintain proliferative cell populations and regenerate robustly (Figs. 4, 6). This reduced cell proliferation in primary tentacle tissue appears to shift towards their adult structure, the anchors, which retain a population of proliferative cells (Fig 3).

### Similar cell dynamics regulate differential investment into juvenile and adult tissues

Most medusozoans make fundamentally different tentacles in their polyp and medusa stages, despite both tissues being used for prey capture. Juvenile tentacles are regressed and new adult tentacles are made during metamorphosis (cubozoans) [10] or post-metamorphosis (scyphozoans) [reviewed by 9]. These tentacles are distinct in their musculature, cnidocyte composition, and cell proliferation patterns [30, 31]; these morphological differences may enable juveniles and adults to capture different prey. Across medusozoans, cell proliferation shows a consistent shift from a wide distribution in polyp tentacles to a restricted distribution in mature medusa tentacles (Fig 8A). Prior to tissue remodeling, proliferative cells can be found throughout the entire juvenile tentacle in *Aurelia sp*. (Scyphozoa) [31], *Tripedalia cystophora* (Cubozoa) and *Alatina moseri* (Cubozoa) [30]. Early medusa tentacles show broad cell proliferation, however, this proliferation is restricted to the base of mature tentacles (*i.e.*, proximal tentacle) in hydrozoans (*Clytia hemisphaerica*, *Cytaeis uchidae*, *Rathkea octopunctata*, and *Cladonema pacificum*) [32, 33], scyphozoans (*Aurelia sp.*) [31], and cubozoans (*T. Cystophora* and *A. moseri*) [30]. It has been hypothesized that the restriction of proliferative cells to the base of adult tentacles may be an ancestral feature of medusozoans [31]. We have observed similar dynamics in *H. sanjuanensis* where growing secondary tentacles exhibit broad cell proliferation and mature secondary tentacles have restricted cell proliferation (Fig 4,8); however, contrary to other medusozoans, cell proliferation is restricted to a collar underneath the tentacle tip, rather than the proximal region of the tentacle. This suggests that the patterns of cell proliferation in adult staurozoan tentacles may be derived. Further, primary tentacles do not show diffuse cell proliferation as seen in the juvenile tentacles of scyphozoans and cubozoans. Instead, they show similar concentrations of proliferative cells below the tentacle tips as found in mature secondary tentacles. (Figs 3,4,8). In addition, previous studies have shown similar musculature and innervation in primary and secondary tentacles [23, 24], suggesting that limited metamorphosis may lead to the re-deployment of a juvenile structure in the adult stage. The shift of cell proliferation patterns during medusozoan tentacle development suggests that cell proliferation may be a mechanism to reduce investment in juvenile tissues. At the onset of tentacle regression, proliferative cells are not detected in the juvenile tentacles of *Aurelia spp*. [34, 35] and *T. cystophora* [30]. Following metamorphosis, cell proliferation is found throughout the re-growing juvenile tentacles of *Aurelia sp.*, indicating that the juvenile tentacle has regained investment [31]. Our results show that reduction of proliferative cells in primary tentacles in *H. sanjuanensis* leads to degeneration of this juvenile structure. Together, these results suggest that primary and secondary tentacles are patterned by a similar developmental program but receive different cues to maintain or reduce their proliferative cell populations. We propose that re-deploying a juvenile tentacle in a different location allows an organism to achieve adult function even in the absence of true metamorphosis.

**Fig 8.**
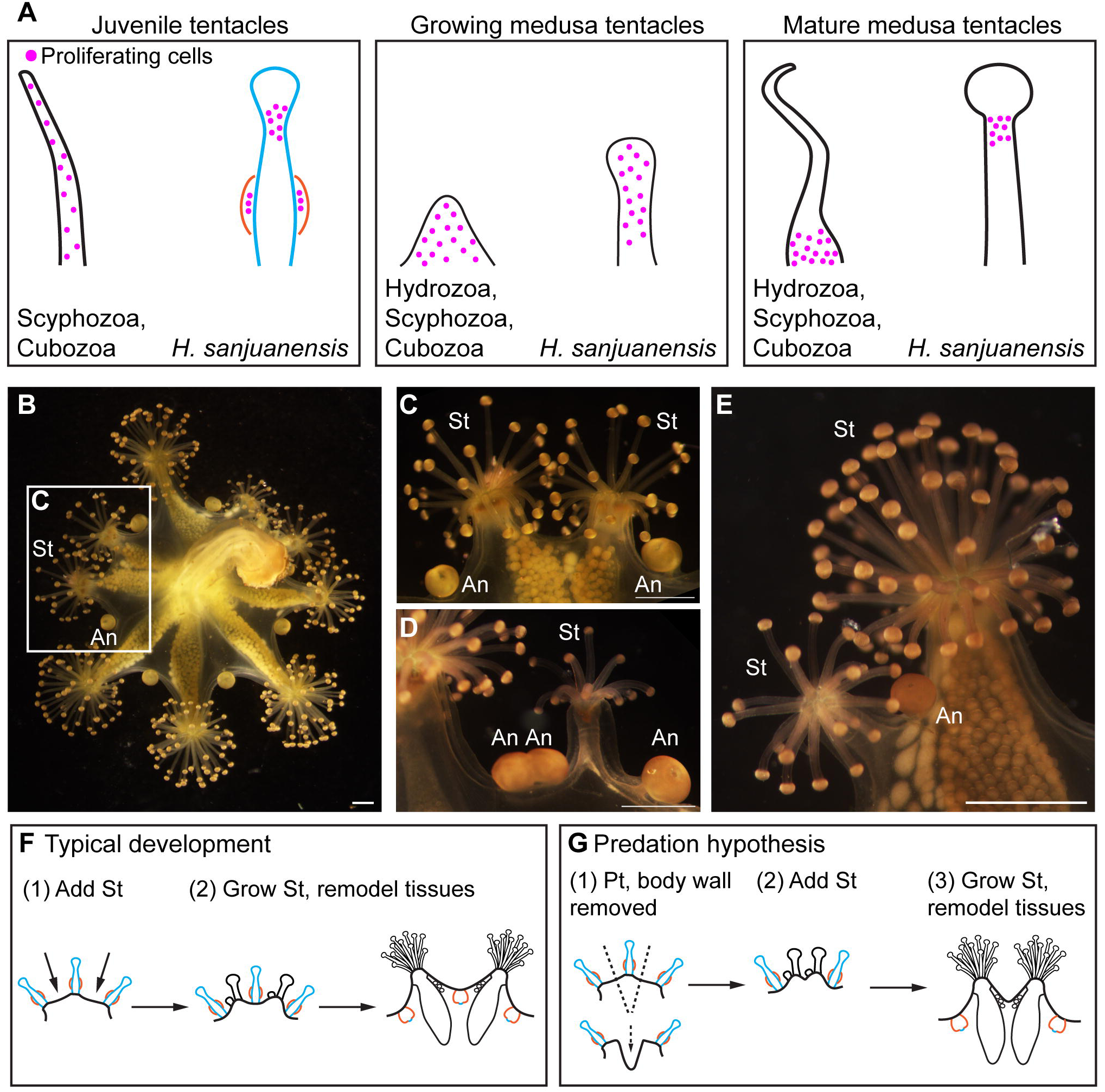
Summary of medusozoan tentacle cell proliferation and abnormal morphologies found in *H. sanjuanensis*. (A) Cell proliferation patterns described for scyphozoans, cubozoans, and hydrozoans and observed for *H. sanjuanensis* in juvenile tentacles (primary tentacles), growing medusa tentacles (secondary tentacles), and mature medusa tentacles (secondary tentacles). Scyphozoan tentacles described for: *Aurelia sp*. [31]. Cubozoan tentacles described for: *Tripedalia cystophora* and *Alatina moseri* [30]. Hydrozoan tentacles described for: *Clytia hemisphaerica*, *Cytaeis uchidae*, *Rathkea octopunctata*, and *Cladonema pacificum* [32, 33]. (B) An example of an adult *H. sanjuanensis* with abnormal morphology. (C) High magnification of the branched secondary tentacle clusters in (B), where two secondary tentacle clusters are present without an anchor between them. (D) A second example of a morphologically abnormal adult animal, where two anchors are joined together and a smaller secondary tentacle cluster sits between the double anchors and the next normal anchor. (E) A third example of a morphologically abnormal adult, where a smaller secondary tentacle cluster and an anchor branch off of a developed secondary tentacle cluster. (F-G) Cartoons summarizing typical development of secondary tentacle clusters and a hypothesized mechanism to generate the morphology observed in (B,C). Scale bars 1 mm (B-E). Anchors (An), secondary tentacles (St).

### Do juvenile tissues scaffold novel adult tissues?

Juvenile tentacles are thought to be homologous across scyphozoans, cubozoans, and staurozoans (primary tentacles) [36 as cited by 17]; both scyphozoans and cubozoans remodel their juvenile tentacles into rhopalia- structures that contain specialized cells and structures to detect light and gravity [10, 37]. As cell proliferation is lost in the juvenile tentacle, a high density of proliferating cells is observed and maintained within these rhopalia [30,35]. We see this same increase in proliferative cells in developing staurozoan anchors (Fig 2), which have a proposed homology to rhopalia [36 as cited by 17; 24, 38]. In other staurozoan lineages, primary tentacles may be maintained, or regressed and not replaced, suggesting that the fate of primary tentacles is heterogeneous across this clade [17]. Maintenance of the primary tentacle across staurozoan lineages, regardless of tentacle fate, suggests there is selection acting to retain this tissue outgrowth. Given the prevalence of remodeling juvenile tentacles, we hypothesize that these juvenile tissues may be important to scaffold the tissues of adult medusozoans. To determine whether primary tentacle development is a necessary precursor for the developing anchor, it will be informative to cut primary tentacles prior to anchor development and determine whether anchors can develop *de novo*, or whether primary tentacles must first develop before the anchor can form. Alternatively, these retained primary tentacles may serve as landmarks, or indicators of position, for the adult body plan, as secondary tentacle clusters develop between these juvenile structures (Fig 2A) [17 and references therein]. Whether primary tentacles are used to scaffold adult anchors, or whether these act as landmarks during metamorphosis suggests an interesting framework with which to test the coupling of juvenile and adult body plans. Examples of coupling between pre- and post-metamorphic body plans have been found in beetles, where disrupting larval leg development impacted adult leg morphology [39] and hornworms, where adult legs are comprised of adult-specific and polymorphic cell populations [40]. Several studies have observed abnormal morphologies in staurozoans, where the incorrect location and number of secondary tentacles and anchors has been observed and suggested to arise from incorrect regeneration due to predation [41–43]. We have additionally observed these abnormalities: including the incorrect number of secondary tentacles found between anchors and duplicated anchors between secondary tentacles (Fig 8B-E). We propose that improper regeneration may be linked to removal of the primary tentacle, prior to metamorphosis, which would eliminate the positional information necessary for specifying the proper location for secondary tentacle development (Fig 8F, G). Explicitly testing this hypothesis will provide an important framework for understanding the role of juvenile structures in dictating the diversity of adult body plans.

## Acknowledgements

This work was supported by the National Institutes of Health (NIGMS R35GM147253-01 to LSB), institutional funds from Cornell University, and funding from Friday Habor Laboratories (Alan J. Kohn and Alexander Fodor Graduate Student Endowed Fellowships to KB). Quantitative image analysis was performed with Imaris Software (Oxford Instruments, UK) at the Cornell Institute of Biotechnology’s BRC Imaging Facility (RRID:SCR_021741). We are grateful to Dr. Claudia Mills for her help with identification of the staurozoan populations on San Juan Island and for her thoughtful comments on this manuscript.

